# A Unified Approach for Identifying PET-based Neuronal Activation and Molecular Connectivity with the functional PET toolbox

**DOI:** 10.1101/2024.11.13.623377

**Authors:** Andreas Hahn, Murray B. Reed, Christian Milz, Pia Falb, Matej Murgaš, Rupert Lanzenberger

**Author notes:** **Correspondence to:** Andreas Hahn, Assoc.Prof. PD PhD MSc, or Rupert Lanzenberger, Prof. PD MD, Department of Psychiatry and Psychotherapy Medical University of Vienna, Austria Waehringer Guertel 18-20, 1090 Vienna, Austria.

## Abstract

**Purpose:** Functional PET (fPET) enables the identification of stimulation-specific changes of various physiological processes (e.g., glucose metabolism, neurotransmitter synthesis) as well as computation of individual molecular connectivity and group-level molecular covariance. However, currently no consistent analysis approach is available for these techniques. We present a versatile, freely available toolbox designed for the analysis of fPET data, thereby filling a gap in the assessment of neuroimaging data.

**Methods:** The fPET toolbox supports analyses for a variety of radiotracers, scanners, experimental protocols, cognitive tasks and species. It includes general linear model (GLM)-based assessment of task-specific effects, percent signal change and absolute quantification, as well as independent component analysis (ICA) for data-driven analyses. Furthermore, it allows computation of molecular connectivity via temporal correlations of PET signals between regions and molecular covariance as between-subject covariance using static images.

**Results:** Toolbox performance was validated by analysis protocols established in previous work. Stimulation-induced changes in [^18^F]FDG metabolic demands and neurotransmitter dynamics obtained with 6-[^18^F]FDOPA and [^11^C]AMT were robustly detected across different cognitive tasks. Molecular connectivity analysis demonstrated metabolic interactions between different networks, whereas group-level covariance analysis highlighted interhemispheric relationships. These results underscore the flexibility of fPET in capturing dynamic molecular processes.

**Conclusions:** The toolbox offers a comprehensive, unified and user-friendly platform for analyzing fPET data across a variety of experimental settings. It provides a reproducible analysis approach, which in turn facilitates sharing of analyses pipelines and comparison across centers to advance the study of brain metabolism and neurotransmitter dynamics in health and disease.

## INTRODUCTION

Identifying neuronal activation in response to specific stimulation is a cornerstone of neuroscience. Blood-oxygen level dependent (BOLD) fMRI is one of the most commonly used techniques and since its discovery in the 1990’s [1] numerous software tools have been made available, such as SPM (https://www.fil.ion.ucl.ac.uk/spm/), FSL (https://fsl.fmrib.ox.ac.uk/fsl/fslwiki/), and AFNI (https://afni.nimh.nih.gov/). While fMRI has greatly increased our understanding of brain function in health and disease, the BOLD signal represents an indirect marker of neuronal activation, comprising relative changes in cerebral blood flow, volume and oxygenation [2, 3].

A more recent approach reflecting neuronal activation is functional positron emission tomography (fPET), which uses radioligands like [^18^F]FDG as a proxy of glucose metabolism [4–6]. Similar to fMRI, fPET employs repeated stimulation and analysis techniques such as the general linear model (GLM) or independent component analysis (ICA) to extract task-specific effects. Since [^18^F]FDG binds almost irreversibly during the scan, a (bolus+) constant infusion protocol is required to maintain free radioligand throughout the experiment, enabling it to bind according to the actual metabolic demands [5]. A major advantage of this technique is the gain in experimental flexibility, allowing several conditions to be imaged in a single scan, whereas conventional bolus application would require separates scans for each condition. The temporal resolution has been gradually improved from minutes to just a few seconds, allowing direct comparison between different signals in the temporal domain [7].

This technique has demonstrated robust increases in glucose metabolism across various cognitive tasks [8–11], with high test-retest reliability [12]. Furthermore, the approach has been successfully extended to image dynamics of neurotransmitter synthesis during task performance, such as the dopamine [13] and serotonin system [14] with 6-[^18^F]FDOPA and [^11^C]AMT, respectively. As fPET provides specific information about physiological processes (such as glucose metabolism, neurotransmitter action, etc.), it offers critical complementary insights into neuronal activation [9, 10, 15], making it particularly valuable to explain effects observed with less specific but commonly used imaging approaches such as BOLD fMRI or EEG. However, until now researchers have relied on own in-house software implementations for fPET data analysis, which pose challenges such as the need for local adaptation and the risk of variability in outcome metrics, leading to reduced reproducibility across studies and imaging centers.

We aim to solve the above issues by providing a versatile toolbox, filling a gap in neuroimaging data analysis. This allows for standardized analyses of fPET data, which facilitates comparison across different tasks, studies and imaging centers as well as sharing of analysis procedures. The current work consolidates a decade of experience with this technique [5, 7–21] into a freely available open-source software package. The toolbox includes Matlab code that has been successfully applied to various data sets involving different stimulation paradigms (visual stimulation, finger tapping, Tetris, working memory, monetary incentive delay, optogenetic stimulation, resting-state) at a range of temporal resolutions (from 60s to 3s). It supports diverse imaging and infusion protocols (constant infusion and bolus + constant infusion), radiotracers ([^18^F]FDG, 6-[^18^F]FDOPA and [^11^C]AMT), scanner systems (GE Advance, Siemens Biograph Vision mMR, 600 Edge and Quadra, Brain Biosciences CerePET, Bruker small-animal PET insert for 7T MRI ClinScan) and species (humans, non-human primates, rodents). The toolbox provides scripting for efficient batch processing and includes a user-friendly graphical user interface (GUI). It will be regularly updated based on new developments and community feedback. This equips basic science and clinical researchers with a generalizable, reproducible analysis approach, which facilitates the widespread use of fPET.

## METHODS

### Implementation

The workflow of the fPET toolbox is shown in figure 1. The process begins by creating an input variable that contains all relevant information for subsequent calculations. This can be done within a Matlab script or automatically via the GUI provided with the toolbox. The input variable ‘fpetbatch’ is a Matlab structure array (struct), which includes details about the process to be executed (e.g. GLM, ICA, molecular connectivity or covariance), input files (e.g., 4D fPET data, masks, blood data, etc.) and further optional parameter settings (e.g., design matrix, filter specifications, etc.). Once the master function is called with the input parameters, all required subfunctions are executed internally without requiring further user input. All settings and intermediate calculations are stored in a file within the designated results directory, facilitating documentation, analyses sharing and troubleshooting.

**Figure 1:**
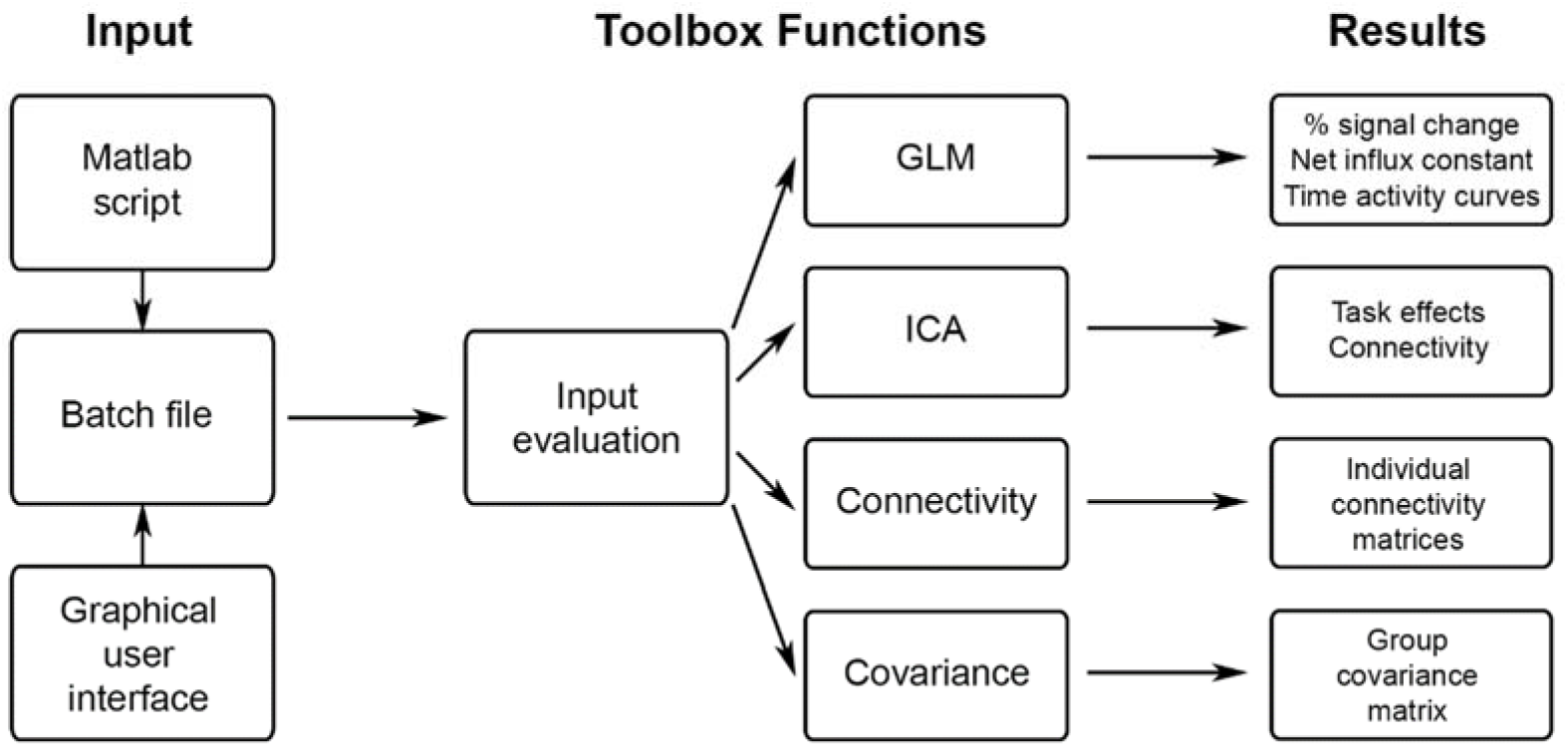
Schematic overview of fPET toolbox functionalities. The input is defined by a script or the GUI and contains all relevant parameters such as the desired calculation, input data, masks, atlases, etc. The toolbox is started by calling a single master function, all other computations are carried out internally, without any further user input. After evaluating the correctness and completeness of the input, the selected calculations are done (GLM, ICA, molecular connectivity or covariance). The corresponding results are then saved in the results folder and can be used for further analyses.

### Functionality

When the toolbox is initiated, it first checks whether data already exists in the specified results directory and provides an option to overwrite the data. It then verifies the completeness and validity of all input variables, alerting the user to any necessary input modifications. Depending on the chosen functionality, calculations are performed on a voxel-wise level for GLM and ICA or region of interest basis for molecular connectivity and covariance.

#### General linear model

The GLM separates stimulation specific effects from baseline by estimating the fit of a predefined model to the data [5, 8]. The design matrix setup is highly flexible with options to define task regressors as ramp functions (slope = 1 kBq/min), custom regressors (e.g., nuisance regressors, non-standard stimulation regressors, etc.) and realignment parameters from motion correction. Baseline radiotracer uptake is defined by an average time activity curve (TAC) as extracted from a user-defined mask (typically gray matter). This baseline can be refined by excluding certain brain areas (e.g. fMRI activation maps, meta-analysis masks, anatomical regions, etc.). Additionally, a third order polynomial function can be used to model the baseline [5], yielding similar stimulation-induced effects [8, 16] and test-retest reliability [12], but less optimal fitting [8].

Before GLM calculations, data can be low-pass filtered and regressors orthogonalized. Solving the GLM (Eq. 1) yields 3D maps of beta values and t-statistics for each regressor (including a baseline term) and residuals.

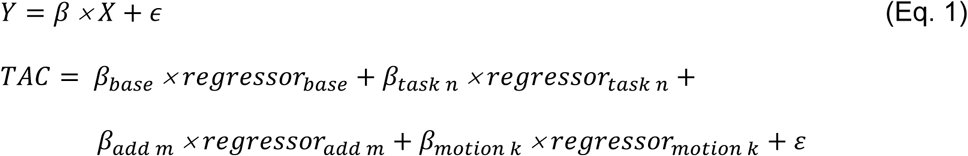

##### Percent signal change

For each stimulation regressor defined in the GLM, 3D maps of percent signal change can be calculated as recently validated [17]. In short, GLM beta estimates of task effects are contrasted with the baseline. This is feasible as under certain assumptions the radiotracer concentration in blood plasma cancels out and percent signal change can be calculated as in equation 2.

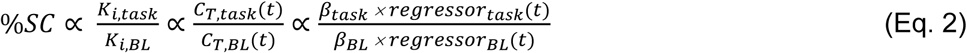

This non-invasive alternative has been shown to yield strong agreement with percent signal change obtained from the absolutely quantified values [17].

##### Absolute quantification

If blood sampling data is available, absolute quantification of the net influx constant Ki can be done for both the baseline and stimulation regressors defined in the GLM using equation 3.

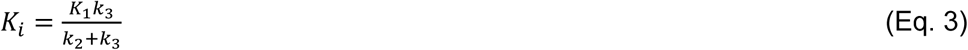

The calculation requires a plasma input function characterizing the concentration of radiotracer in blood or to compute this via a whole-blood input function and plasma-to-whole blood ratio. Blood data are linearly interpolated to match the PET data frame times. The quantification is then carried out with the Patlak plot [22] (Eq. 4), where the start of the linear fit is set to t* = 1/3 of the scan duration.

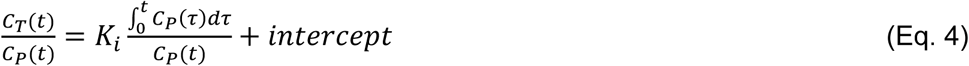

For [^18^F]FDG the cerebral metabolic rate of glucose (CMRGlu) can be calculated (Eq. 5) if blood glucose levels (Glu_P_) are provided, using a default lumped constant of LC = 0.89 [23, 24].

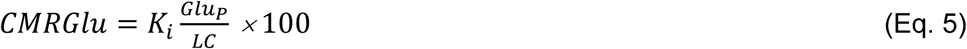

##### Visualizing task effects

In addition to voxel-wise maps of outcome parameters, the temporal characteristics of stimulation effects can be visualized. This function isolates the task-specific TAC for a particular brain region (as defined by a mask image) by subtracting all non-stimulation related parameters of the GLM (see Eq. 1). Visualization can be performed at both, the individual and group level.

#### Independent component analysis

fPET data can also be analyzed using ICA, a data-driven alternative to the GLM that identifies stimulation-specific effects without prior knowledge of their timing or spatial extent. The ICA approach is model-free, reducing the initial setup complexity but requiring careful interpretation of the results. Both, task-specific and resting-state fPET data can be processed.

The ICA implementation uses FastICA [25], which is widely used for analysis of e.g., BOLD fMRI data. However, since radiotracers currently used for fPET exhibit irreversible kinetics, it is required to remove the baseline uptake beforehand [15, 26]. This is achieved by z-scoring each time frame, after dimensionality reduction via principal component analysis. Depending on the number of input files, time courses of each individual are concatenated to enable group ICA analysis. Since FastICA uses random starting points, repeated use may lead to different results. To provide stable component estimates, FastICA is run ten times internally. For each run, the kurtosis of ICs is computed [15] and the solution with the highest kurtosis (average of the top 50% of ICs) is selected.

#### Molecular connectivity

Molecular connectivity assesses the similarity between PET signals across brain regions based on their molecular characteristics. We would like to emphasize that the terminology is employed according to a recent consensus within the field [20], which has also been employed by other groups [27]. In essence, the term ‘connectivity’ describes the relationship between brain regions based on the similarity of their signal’s temporal characteristics, like in fMRI and EEG research. As such, this is calculated within a single individual. In contrast, the term ‘covariance’ refers to the calculation of associations between groups of subjects.

Similar to ICA, this requires removal of the baseline radiotracer uptake, which can be achieved by fitting each ROI’s TAC with a third order polynomial, regression against a representative baseline TAC or spatio-temporal filtering [19, 28]. An ROI atlas is used to compute pairwise correlations between regions, resulting in a molecular connectivity matrix for each individual.

#### Molecular covariance

Molecular covariance refers to the calculation of associations between subjects using static 3D maps (SUV, Ki, volume of distribution, binding potential, etc.) and an ROI atlas to calculate pairwise regional correlations. This type of analysis already began over 40 years ago [29, 30]. Unlike molecular connectivity (which uses temporal information), covariance is computed across subjects rather than within an individual, enabling to draw inferences at the group level only. For more detailed discussions about this and related issues see other recent work [20, 28, 31, 32]. For the computation of molecular covariance, an intensity normalization may be required to remove spurious intersubject variability [33, 34].

### Environment

The free and open-source toolbox is written in Matlab and tested on versions R2021b update 3 and R2018a, It relies on SPM12 for data handling (https://www.fil.ion.ucl.ac.uk/spm/) and the FastICA toolbox v2.5 (http://research.ics.aalto.fi/ica/fastica/index.shtml), both of which are distributed under the GNU General Public License.

### License and disclaimer

The toolbox is and will be actively maintained by the authors through official channels, but also accepts contributions from the research community.

The fPET toolbox is open-source software released under the terms of the GNU General Public License version 2 as published by the Free Software Foundation (https://www.gnu.org/licenses/old-licenses/gpl-2.0.html). In short, it may be modified and distributed for non-commercial purposes, provided the original source is cited. The software is provided ’as is,’ without any warranty, express or implied, including but not limited to warranties of fitness for a particular purpose. The developers shall not be held liable for any damages or issues arising from its use.

## RESULTS

The toolbox’s functionality is demonstrated using previously published human data. This includes visual stimulation and finger tapping (n=5) [8] and the video game Tetris (n=24) [11], both scanned with [^18^F]FDG. Data from 6-[^18^F]FDOPA (n=10) [18] and [^11^C]AMT (n=16) [14] were obtained while subjects performed the monetary incentive delay reward task. Molecular connectivity and covariance were computed using resting-state [^18^F]FDG bolus + infusion data from n=8 subjects [19]. An extensive description of the experimental design, imaging protocols and demographic information can be found within the respective manuscripts. Application of the toolbox effectively reproduced the previously reported findings across various settings, including different radiotracers, scanner systems and cognitive tasks.

The GLM analysis separated task effects from baseline radiotracer uptake and absolute quantification with arterial input functions yielded CMRGlu and Ki. Data obtained with [^18^F]FDG while subjects completed the Tetris task yielded significant activation in the intraparietal sulcus, frontal eye field and visual cortex (Fig. 2a) [10, 11, 16], with percent signal changes of approximately 20% from baseline. Likewise, the monetary incentive delay task showed increases in 6-[^18^F]FDOPA dopamine synthesis rates in the ventral striatum (Fig. 2b) [18] as well as [^11^C]AMT serotonin synthesis rates in the ventral striatum and anterior insula (Fig. 2c) [14]. Here, task-induced changes reached approximately up to 150% and 40% for the two neurotransmitter systems, respectively. Evaluation of task-specific time activity curves confirmed the model fits in the temporal domain for each of the different radiotracers.

**Figure 2:**
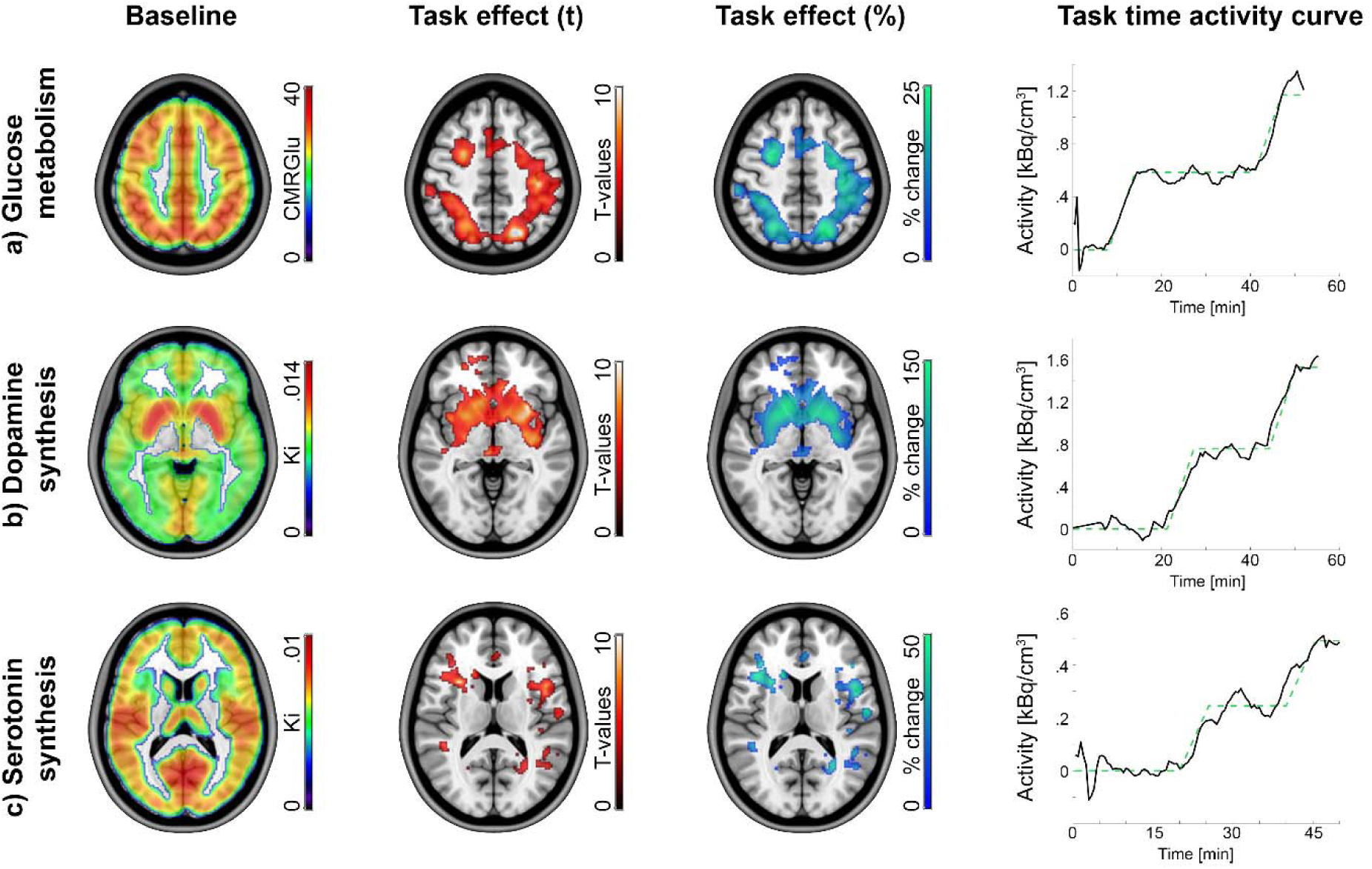
Exemplary results obtained with the GLM. The approach separates baseline radiotracer uptake (column 1) from task-specific effects (columns 2-4) as defined by the design matrix. Quantification was done in an absolute manner with the Patlak plot (CMRGlu, Ki) and relative to baseline as percent signal change (%). Task-specific effects were subject to a one sample t-test (p<0.05 FWE corrected cluster level, following p<0.001 uncorrected voxel level). Task-specific time activity curves were extracted to visualize model fits. The cerebral metabolic rate of glucose (CMRGlu, **a**) was obtained with [^18^F]FDG, while subjects completed the Video game Tetris® [10, 11, 16]. The net influx constant Ki of dopamine (**b**) and serotonin synthesis rates (**c**) was obtained with 6-[^18^F]FDOPA [18] and [^11^C]AMT [14], respectively, while subjects carried out the monetary incentive delay reward task.

ICA of [^18^F]FDG data during visual stimulation and right finger tapping [8] identified the corresponding component of task-specific activation. This included the primary visual cortex as well as the primary motor cortex and supplementary motor area (Fig. 3).

**Figure 3:**
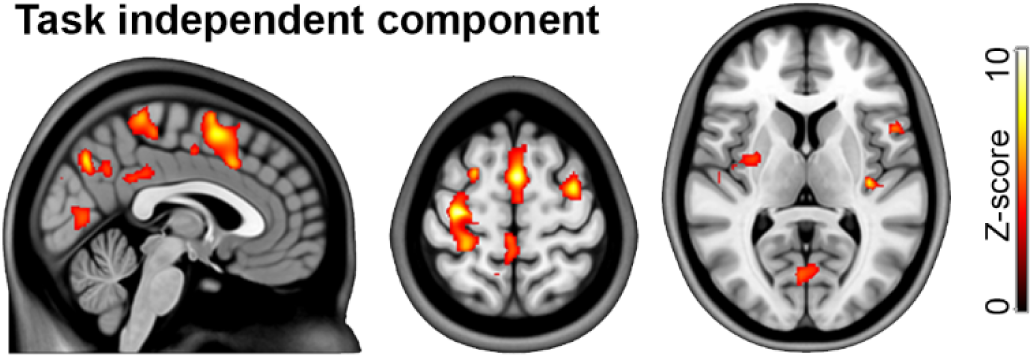
Exemplary results obtained with ICA. After removal of baseline radiotracer uptake, the approach isolates independent components based on similar spatio-temporal characteristics. Task effects were obtained with [^18^F]FDG, while subjects underwent visual stimulation and simultaneously performed a right finger tapping task [8].

Computation of molecular connectivity and covariance from resting-state [^18^F]FDG scans was done with dynamic fPET data (50 x 1min frames) and static images thereof (average of last 10 min), respectively [19]. As demonstrated previously [19, 28], within-subject molecular connectivity and between-subject covariance yielded different network patterns. Molecular connectivity was characterized by strong links between different modules, while covariance showed strongest links between homologous brain regions (Fig. 4a and b).

**Figure 4:**
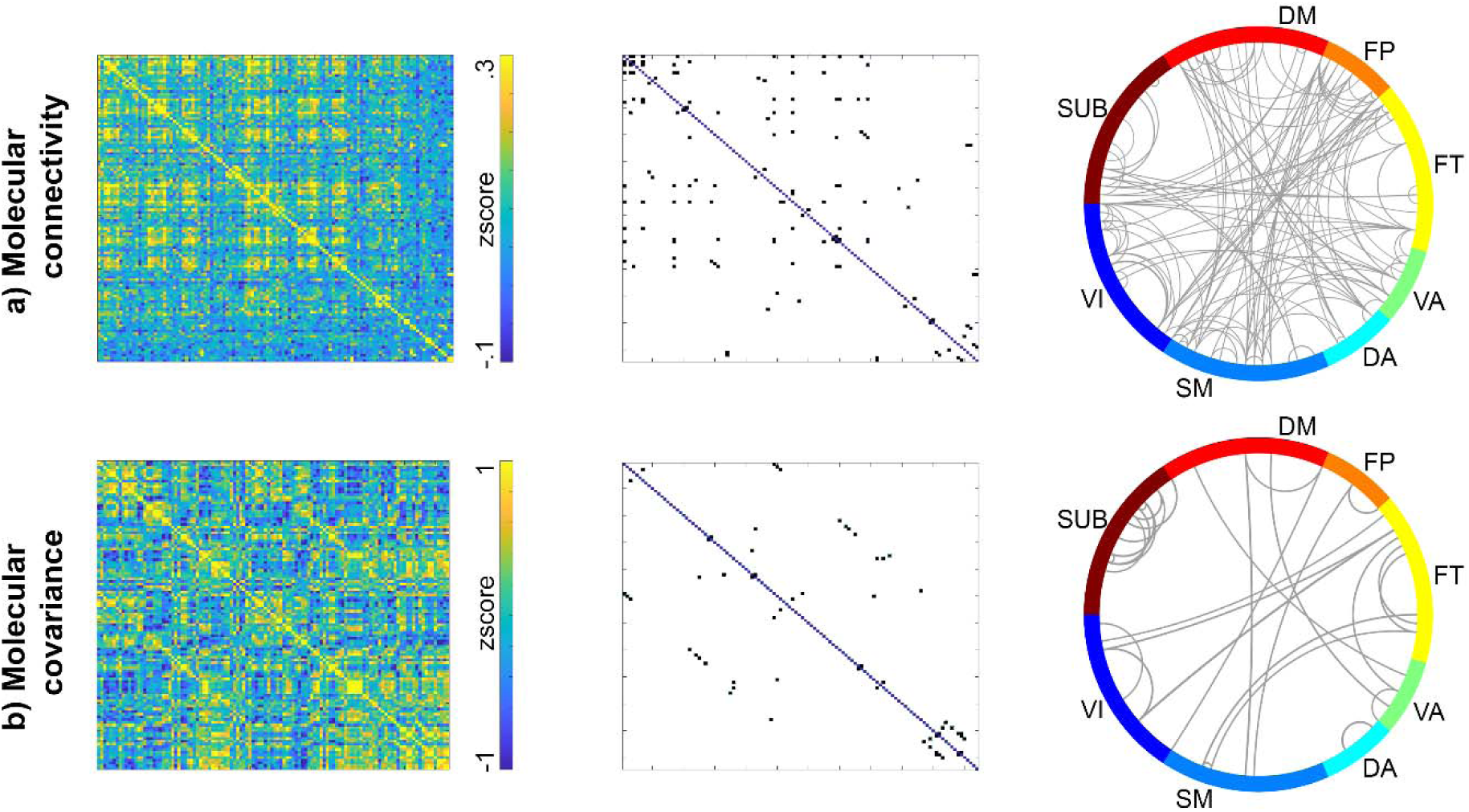
Exemplary results of molecular connectivity and covariance. **a)** For molecular connectivity, the baseline was removed by fitting a third-order polynomial to each region’s TAC. Pairwise regional correlations were calculated for each individual as temporal correlations. **b)** Molecular covariance was computed from static images as correlation between subjects. Resting-state fPET data for both analysis were acquired with [^18^F]FDG [19]. For a) the full dynamic dataset was included, while for b) a static image was created as mean of the last 10 min. Cortical and subcortical regions were taken from the Harvard-Oxford atlas (left and right hemisphere separated). Thresholded matrices were obtained by retaining connections, which are

## DISCUSSION

We introduce a versatile toolbox for the analysis of fPET data, enabling the identification of stimulation-specific changes across various physiological processes, including glucose metabolism as well as dopamine and serotonin synthesis. In addition to task-based analyses, the software also supports computation of individual molecular connectivity and group-level covariance matrices with virtually any radioligand. The centralization of all parameter settings in a single file provides an effective approach to document, share and troubleshoot analysis workflows.

### Methodological considerations

Although this work does not aim to provide a comprehensive comparison between all available methodologies, a few advantages and limitations are noteworthy. The GLM approach is particularly suitable for situations where a clear hypothesis exists regarding the timing of activation, often the case in task-based studies [35, 36]. This approach enables robust detection of task-specific effects with intuitive interpretability due to the pre-defined model. Furthermore, GLM estimates can be extended to derive percent signal change or perform absolute quantification of the net influx constant [5, 17]. However, the GLM effectiveness depends on how well the model matches the actual data. Underestimation of effects may occur if unexpected or complex temporal variations occur. In contrast, ICA is data-driven and does not require a priori knowledge on the timing and shape of the effects, making it suitable for detecting spontaneous or unknown activity sources that exhibit more complex signals [6, 26]. ICA is also advantageous for noise removal [37], particularly for high-temporal resolution data where physiological artefacts such as respiratory or cardiac signals, can become significant [38]. However, ICA has its limitations. The components can be biologically challenging to interpret, particularly when the data contains mixed sources or artefacts and it may show lower sensitivity and specificity to task-relevant activation, due to its model-free nature. Furthermore, the selection of the number of components can be subjective, requiring validation [39].

The fPET toolbox supports both individual molecular connectivity and group-level molecular covariance analyses, each offering distinct insights. Molecular connectivity captures the temporal similarity of radiotracer uptake between brain regions within an individual, akin to functional connectivity in fMRI. This is particularly useful for studying network interactions based on metabolic demands and neurotransmitter systems like dopamine and serotonin, revealing synchronized activity patterns in response to tasks or pharmacological challenges. However, interpreting such connections still requires further work to as this is not yet established beyond [^18^F]FDG. On the other hand, group-level covariance aggregates data across individuals to explore consistent patterns of static molecular characteristics. This approach is more suitable for population-level studies, such as comparing patient groups or examining broad network disruptions. Covariance analysis highlights how molecular processes differ across subjects, providing a macroscopic view of brain function. Thus, the two methods offer complementary insights. Molecular connectivity is ideal for individualized, real-time analysis, while covariance excels in population studies or situations where no dynamic data is available. The toolbox allows for flexible application of both approaches, enabling tailored analyses depending on the research focus.

### Recommendations for future studies

To stimulate future research with fPET, we provide several key recommendations for its optimization (Table 1). While early fPET studies utilized a constant radioligand infusion [4, 5], the bolus + constant infusion protocol offers clear advantages. This infusion protocol increases the signal-to-noise ratio, particularly at early time points [8], thereby improving motion correction accuracy. Moreover, it generates TACs with near-linear characteristics, enabling more robust modelling and reduced bias task effect estimates [5, 8, 40]. Although the optimal ratio between bolus and subsequent infusion has yet to be identified [8, 40], higher infusion rates lead to more radioactivity for the estimation of task effects and thus more robust task estimation, due to a steeper slope of the TAC. This aligns with the earliest work on task-specific changes in metabolic demands, which applied the stimulus during the initial bolus [41], coinciding with the TAC’s steepest slope.

**Table 1:** Recommended experimental settings for fPET studies. These settings have been successfully applied across various analysis techniques (GLM, ICA, molecular connectivity and covariance), condition (rest or task), radiotracer ([^18^F]FDG, 6-[^18^F]FDOPA, [^11^C]AMT), species (humans, non-human primates, rodents) and PET scanners. Nevertheless, we acknowledge that special applications and further advancements may re-define the optimal settings.

| Aspect | Recommendation | Reason |
| --- | --- | --- |
| Radiotracer administration | Bolus + constant infusion | Enhances overall signal-to-noise ratio (SNR) and produces nearly linear time-activity curves (TACs), enabling more accurate motion correction and model robustness. |
|  | Bolus activity = 20% | Ensures that the majority (80%) of radiotracer activity is constantly infused throughout the measurement and is thus readily available for stimulation-induced effects. |
| Task design | Initial rest period ~ 6-8 min | Allows for stabilization of radiotracer uptake, establishing a reliable baseline for subsequent task-induced signal changes. |
|  | Rest periods between tasks | Aids in isolating task-related changes by differentiating them from baseline activity, improving signal attribution. |
|  | Final rest period | Helps distinguishing task-related signal changes at the end of the scan, reducing ambiguity in identifying post-task activity. |
|  | Similar length for task and rest | Balances task and rest durations, which can lead to more consistent task responses and more reliable signal detection. |
| Blood sampling | During rest periods | Minimizes interference with task engagement, reducing movement-related artifacts and ensuring clearer signal attribution to task vs. baseline. |
| GLM analysis | Calculate percent signal change | Provides a standard, non-invasive metric for comparing activation across studies, facilitates reproducibility, eliminates need for blood sampling without extra user input. |
| ICA analysis | Remove initial rest period | Avoids contamination of ICA by initial frames with low SNR, ensuring more accurate task-related component extraction. |
| Molecular connectivity analysis | Remove initial rest period | Prevents initial uptake fluctuations from skewing connectivity estimates, leading to more reliable connectivity patterns. |
| Molecular covariance analysis | Appropriate normalization mask | Reduces intersubject variability by accounting for individual differences in uptake, allowing for more robust cross-subject comparisons. |

We further recommend delaying the onset of stimulation until a stable baseline with near-linear uptake is established, approximately 6 min after start of the radiotracer administration. Incorporating rest periods between task blocks, similar to fMRI protocols, aids in distinguishing task-related changes from baseline activity. The inclusion of a rest period following the final task block also helps to avoid ambiguity in task-related effects at the end of the scan. While consensus on such resting periods in functional imaging is limited, a similar length as the stimulation block has been shown to yield robust results, e.g., [16].

For simultaneous PET/MR imaging, the choice between a hierarchical [6] or continuous task design [16] depends on the actual experiment and aim of the study. As a hierarchical design contains short task (and rest) blocks for fMRI within long task blocks for fPET, this might be well suited for simultaneous assessment of neuronal activation proxies obtained with fMRI and fPET. However, one needs to be aware that changes in the [^18^F]FDG fPET signal do indeed occur within seconds, and thus also short task and rest blocks will influence the signal [7]. On the other hand, a continuous design employs continuous task performance for longer time periods of e.g., 5 min, without resting periods during that time. This is particularly suited for the simultaneous assessment of functional connectivity [11, 16] or other MRI techniques like arterial spin labelling or spectroscopy.

When employing the GLM for the identification of task effects, we recommend always calculating percent signal change. This non-invasive measure offers a comparable and scalable outcome across different experiments, radiotracers and research centres, without the need for invasive blood sampling or extra user input. Additionally, plotting and inspecting TACs remains a simple and effective means to inspect data quality, evaluate model fits and detect potential artefacts in the data or potential issues with radioligand infusion.

Finally, the authors offer widespread support for fPET studies, including experimental design, measurement protocol, practical implementation, data analysis and multimodal combinations, based on various international collaborations. Examples of these include previous work on the combination of metabolic demands and functional connectivity [16], the divergence between glucose metabolism and the BOLD signal [9, 10, 15], optimization of the preprocessing pipeline [42] and measurement protocols [18].

### Limitations, outlook and conclusions

Currently, the toolbox focuses on deriving task estimates or connectivity for individual subjects as well as covariance across groups. However, it does not implement statistical testing. The decision is deliberate, as the scope of this work is to provide individual-level outputs that can then be further processed. These can be realized using well-established external tools for group-level statistics, such as SPM, FSL and AFNI for task effects as well as network based statistics [43] and the brain connectivity toolbox [44] for molecular connectivity.

With a decade of international experience, the unified approach offered by this toolbox aims to stimulate further advancements in the field of fPET. This may be particularly relevant for imaging the dynamics of neurotransmitter synthesis, showing unprecedented stimulation-induced signal changes around 100% from baseline with 6-[^18^F]FDOPA [13] and 40% for [^11^C]AMT [14], even for cognitive tasks. Looking forward, fPET holds considerable promise for the identification of pharmacological effects. Particularly, targeting and imaging the same neurotransmitter system may provide insights into the mode of action of neuropsychopharmacological treatment agents and drugs of abuse. These approaches could be further extended to patient populations, where the exploitation of functional dynamics of molecular processes offers a novel view on brain disorders as compared to conventional techniques [45]. Here, the availability of percent signal change as a quantitative metric makes this technique more accessible by eliminating the need for invasive blood sampling.

## ACKNOWLEDGEMENTS

We thank the graduated team members and the diploma students of the Neuroimaging Lab (NIL, head: R. Lanzenberger) as well as the clinical colleagues from the Department of Psychiatry and Psychotherapy for clinical and/or administrative support. In detail, we would like to thank S. Kasper, K. Papageorgiou, G.M. Godbersen, S. Klug, P. Michenthaler, T. Vanicek, A. Basaran, M. Hienert, L. Silberbauer, J. Unterholzner and G. Gryglewski for their medical support, L. Rischka, and M. Klöbl for acquisition and analysis support, V. Ritter, K. Einenkel, E. Sittenberger and D. Gomola for subject recruitment. We are further grateful to L. Nics, C. Vraka, C. Philippe, V. Pichler, J. Raitanen, J. Völkle, A. Pomberger and L. Aichinger for radioligand synthesis. The scientific project was performed with the support of the Medical Imaging Cluster of the Medical University of Vienna.

This research was funded in whole or in part by the Austrian Science Fund (FWF) [Grant DOI 10.55776/KLI610 and 10.55776/KLI1151, PI: A. Hahn, Grant DOI 10.55776/KLI1006, PI: R. Lanzenberger] and the Vienna Science and Technology Fund (WWTF) [10.47379/CS18039], Co-PI: R. Lanzenberger. For the purpose of open access, the author has applied a CC BY public copyright license to any Author Accepted Manuscript version arising from this submission. Christian Milz is a recipient of a DOC Fellowship (27221) from the Austrian Academy of Sciences at the Department of Psychiatry and Psychotherapy, Medical University of Vienna.

## AUTHOR CONTRIBUTIONS

Study design: A.H., M.B.R., R.L.

Methods: A.H., M.B.R.

Data analysis: A.H., M.B.R.

Manuscript preparation: A.H., M.B.R., C.M., P.F., M.M.

All authors discussed the implications of the findings and approved the final version of the manuscript.

## CODE AVAILABILITY

All code and documentation for the fPET toolbox is available on GitHub and is distributed under the GNU general public license version 2.0. https://github.com/NeuroimagingLabsMUV/fPET-toolbox.git

